# Transcranial ultrasound selectively biases decision-making in primates

**DOI:** 10.1101/486134

**Authors:** Jan Kubanek, Julian Brown, Patrick Ye, Kim Butts Pauly, Tirin Moore, William Newsome

## Abstract

Transcranial focused ultrasound has the promise to evolve into a transformative noninvasive way to modulate activity of neuronal circuits deep in the brain. The approach may enable systematic and causal mapping of how individual brain circuits are involved in specific behaviors and behavioral disorders. Previous studies demonstrated neuromodulatory potential, but the effect polarity, size, and spatial specificity have been difficult to assess. Here, we engaged non-human primates (*macaca mulatta*) in an established task that provides a well defined framework to characterize the neuromodulatory effects. In this task, subjects decide whether to look at a right or a left target, guided by one the targets appearing first. Previous studies showed that excitation/inhibition of oculomotor circuits leads to contralateral/ipsilateral biases in this choice behavior. We found that brief, low-intensity ultrasound stimuli (300 ms, 0.6 MPa, 270 kHz) delivered to the animals’ left/right frontal eye fields bias the animals’ decisions to the right/left visual hemifield. The effect was modest, about on the order of that produced when injecting moderate amounts of potent neuromodulatory drugs into the same regions in this task. The polarity of the effects suggested a neuronal excitation within the stimulated regions. No effects were observed when we applied the same stimuli to control brain regions not involved in oculomotor target selection. Together, using an established paradigm, we found that transcranial ultrasound is capable of modulating neurons to the extent of biasing choice behavior of non-human primates. A demonstration of tangible, brain-region-specific effects on behavior of primates constitutes a critical step toward applying this noninvasive neuromodulation method in investigations of how specific neural circuits are involved in specific behaviors or disease signs.

## Introduction

Noninvasive and spatially specific approaches to modulating neuronal activity have the potential to revolutionize the diagnoses and treatments of a variety of brain disorders. One such approach, ultrasound, can be applied through the intact skull and skin and focused into tight regions deep in the human brain (Ghanouni et al., 2015). At the focus, ultrasound has been shown to modulate neural activity (Naor et al., 2016; Fini and Tyler, 2017; Kubanek, 2018; Tyler et al., 2018; Fomenko et al., 2018). By virtue of its noninvasiveness and spatial focus, the approach has a unique promise in modulating the activity of specific circuits in a systematic fashion. This could enable us, for the first time, to characterize the *causal* contribution of specific brain circuits to specific behaviors or behavioral disorders *in humans.* To bring the approach into clinics, we need to be able to establish three critical aspects of the neuromodulatory effects. First, it is crucial to characterize the polarity of the neuromodulatory effects—whether neurons are excited or inhibited by ultrasound. Second, we need to determine how strong the effects are. If the method is to be useful for mapping brain function in a causal manner, the effects on neurons must be strong enough to manifest in behavior. For example, if clinicians are to determine which brain nuclei underlie a patient’s essential tremor, the neuromodulatory effects on a particular nucleus in question must be strong enough to yield measurable changes in the tremor amplitude. And third, it is critical to validate that the neuromodulatory effects are confined to the focal region of the ultrasound.

This information has been difficult to infer from the approaches and metrics used in previous studies. The bulk of work on ultrasonic neuromodulation has been performed in lightly anesthetized mice (Tufail et al., 2011; King et al., 2013; Kim et al., 2014; Mehić et al., 2014; Ye et al., 2016; Kamimura et al., 2016; Li et al., 2016; Sato et al., 2018). In these studies, applications of low-intensity stimuli to peri-motor regions often lead to visible movements of the limbs or other body parts. There have been concerns that the small size of the rodent brain, relative to the dimensions of the focal spot, results in reflections, standing waves, and consequently, artifactual effects (Guo et al., 2018; Sato et al., 2018). Within this debate, it has been argued that the neuromodulatory effects of ultrasound might be local (King et al., 2014; Mehić et al., 2014; Kamimura et al., 2016), but it has also been argued that these effects might be merely due to auditory or vestibular artifacts (Guo et al., 2018; Sato et al., 2018). Either way, the probability of eliciting movements on a given stimulation trial (“success rate”) has been difficult to reproduce consistently and is strongly dependent on the kind and level of anesthesia (Naor et al., 2016). On the other hand, studies using larger mammals including sheep, macaques, and humans have shown effects on aggregate metrics including EEG activity, MRI BOLD, or reaction time (Deffieux et al., 2013; Hameroff et al., 2013; Legon et al., 2014; Lee et al., 2015; Lee et al., 2016; Lee et al., 2016; Wattiez et al., 2017; Legon et al., 2018). It has been difficult to judge from these studies how strong the effects are and in what direction they point because there has been no particular prediction framework within which to interpret these effects. In addition, it has not been clear how local the effects are (Lee et al., 2016).

Here, we characterize the size and polarity of the neuromodulatory effects of ultrasound using a well-established task (Oppenheim, 1885; Rorden et al., 1997; Ro et al., 2001; Schiller and Tehovnik, 2003; Kubanek et al., 2015) in awake behaving non-human primates (NHPs). In this task, a subject decides whether to look at a right or a left target, guided by one of the targets appearing slightly earlier than the other target. Previous studies using pharmacological or electrical interventions showed that specific, neuroinhibitory or neuroexcitatory perturbations of visuomotor regions produce predictable shifts in subjects’ decisions regarding which target to choose. Thus, this prior research provides predictions of what behavior to expect if a neuromodulatory approach such as ultrasound is excitatory or inhibitory. Moreover, these studies enable us to gauge the size of the neuromodulatory effects from the magnitude of the behavior shift. Finally, the large brain of NHPs and hemispheric symmetry allows us to assess whether the neuromodulatory effects are specific to the stimulated regions.

## Results

We engaged two macaque monkeys in a task that is often used in neurology to diagnose the impact of brain lesions such as those induced by stroke (Oppenheim, 1885; Rorden et al., 1997; Ro et al., 2001). In this paradigm (Fig. 1A), one visual target is shown in the left and one in the right visual hemifield, with a short, controlled delay between the onsets. Typically, in this task, healthy, normal subjects tend to look at the target that appeared first. Stroke or lesions of specific nodes of the oculomotor network, such as the frontal eye fields (FEF) or the lateral intraparietal area (LIP), strongly affect this behavior (Rorden et al., 1997; Ro et al., 2001; Schiller and Tehovnik, 2003; Wardak et al., 2004; Kubanek et al., 2015). These oculomotor circuits preferentially represent targets in their contralateral visual hemifield (Fig. 1B). As a consequence, when neurons in these circuits are affected by a stroke, the contralateral visual hemifield is underrepresented, and subjects preferentially decide to look at the ipsilesional target (Rorden et al., 1997; Ro et al., 2001). These effects are also observed for other kinds of neural perturbations, such as when neuromodulatory agents are injected into these regions. For example, when muscimol—a potent neuroinhibitory drug—is injected into left FEF of macaques (Schiller and Tehovnik, 2003), animals performing this task exhibit a strong ipsilateral—leftward bias (Fig. 1C, red). In contrast, injecting a drug with opposite, disinhibitory effects such as bicuculine into the same region (Schiller and Tehovnik, 2003) leads to an opposite, contralateral bias (Fig. 1C, blue). Analogous results are obtained for neuromodulatory interventions into area LIP (Hanks et al., 2006; Schiller and Tehovnik, 2003; Wardak et al., 2004; Kubanek et al., 2015). This task therefore provides a well established framework that enables us to interpret the neuromodulatory effects of ultrasound—or any other intervention, for that matter. Excitatory interventions or stimuli bias subjects’ decisions in the contralateral direction, whereas inhibitory interventions in the opposite direction.

**Figure 1.**
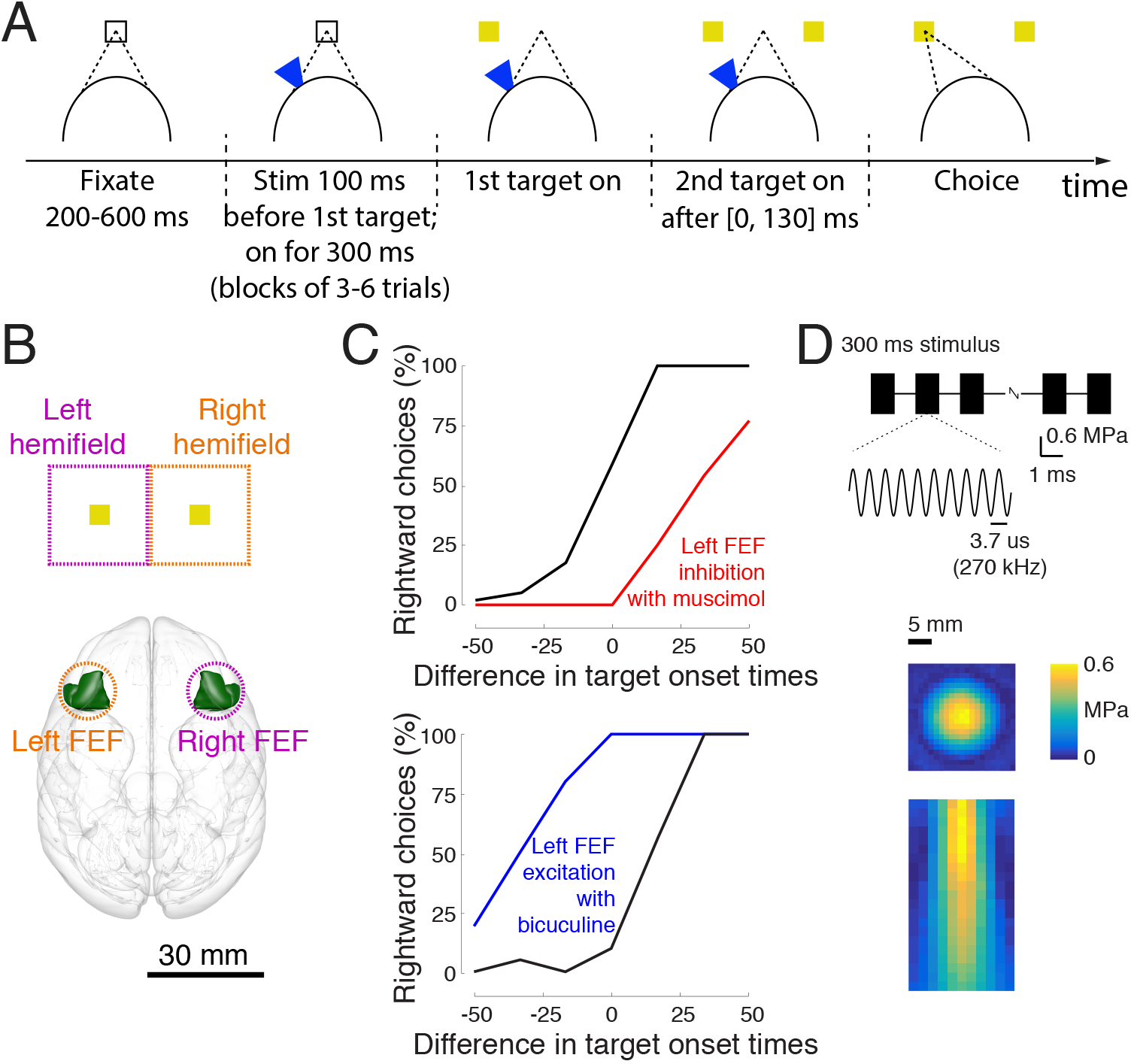
Task and Stimulus. A) Task. The subject fixates a central target. One target appears in the left or the right visual hemifield. After a brief random delay, a second target appears in the opposite hemifield. The subject is free to look at either target after the first target has been presented, and receives a liquid reward if he looks at a target within a 2 ° acceptance window. Ultrasound is applied in blocks of 3-6 trials, strictly interleaved with no stimulation blocks of the same duration, 100 ms prior to the appearance of the first target. B) Functional characterization of the visuomotor system. We delivered the ultrasound non-invasively (intact skull and skin) into the frontal eye fields (FEF). From anatomical and functional studies, it is known that left/right FEF preferentially represents targets in the right/left visual hemifield. The outline of the FEF was rendered using the Calabrese et al. (2015) atlas with Paxinos brain regions. C) Effects in previous studies. When a large amount of strong inhibitory/disinhibitory drugs is injected into left FEF in this task, animals show a strong ipsilateral/contralateral bias in this task (reproduced from Schiller and Tehovnik, 2003, with permission). These results are as expected given the contralateral nature of the visual hemifield representation (B), and are analogous when other nodes of the visuomotor network, such as the parietal area LIP, are perturbed. D) Stimulus. The ultrasound stimulus (0.6 MPa, 270 kHz, 300 ms duration) was pulsed at 500 Hz with 1 ms tone burst duration. The ultrasound was applied through a coupling cone filled with agar gel. The resulting pressure, measured in free field, is provided along the lateral (1 mm steps) and axial (2 mm steps) dimensions in color.

We used this framework to evaluate the polarity and size of the effects of ultrasound on neurons (Fig. 1A). In a given session, ultrasound was applied to the animals’ left or right FEF. Ultrasound was applied in blocks of 3-6 trials and was strictly interleaved with blocks of 3-6 trials in which ultrasound was not applied to exclude potential effects of session time. The stimulus was applied while an animal was making his decision: 100 ms prior to the onset of the first target, and the stimulation lasted for 300 ms. Given that animals responded on average within 171 and 263 ms (monkey A and B, respectively), this perturbation influences a substantial portion of the decision-making process.

The ultrasound stimulus (Fig. 1D) had neuromodulatory parameters (0.6 MegaPa, 270 kHz, 300 ms duration, 500 Hz pulse repetition frequency, 50% duty cycle) that fell into commonly used ranges. We chose a relatively low carrier frequency of 270 kHz so that we could be confident that the stimulus would effectively penetrate the animals’ skull (Deffieux et al., 2013). As a consequence of this choice, the stimulus was relatively broad as indicated by our measurements of free field pressure (Fig. 1D). Full width at half maximum pressure was 10.5 mm in the lateral dimension and 21.8 mm in the axial direction below the skull.

Animals showed typical choice behavior in this task, being sensitive to the difference in target onset times (Fig. 2A, black curves). As in previous studies (Rorden et al., 1997; Ro et al., 2001; Schiller and Tehovnik, 2003; Wardak et al., 2004; Kubanek et al., 2015), the earlier a target appeared before the subsequent target, the more likely the animal was to choose that target. Critically, transcranial ultrasound had a strong influence on this choice behavior (Fig. 2A, blue curves). In the trials in which ultrasound was applied to left FEF (left column), animals were more likely to choose the rightward target. The effect reversed polarity when ultrasound was applied to right FEF (right column): in this case, animals were more likely to choose the leftward target. These single-session effects were significant (*p* < 0.0017, two-tailed two-sample proportion tests) with the exception of the monkey A left FEF example (p = 0.074). These examples suggest that ultrasound biases the animals’ decisions in the contralateral direction.

**Figure 2.**
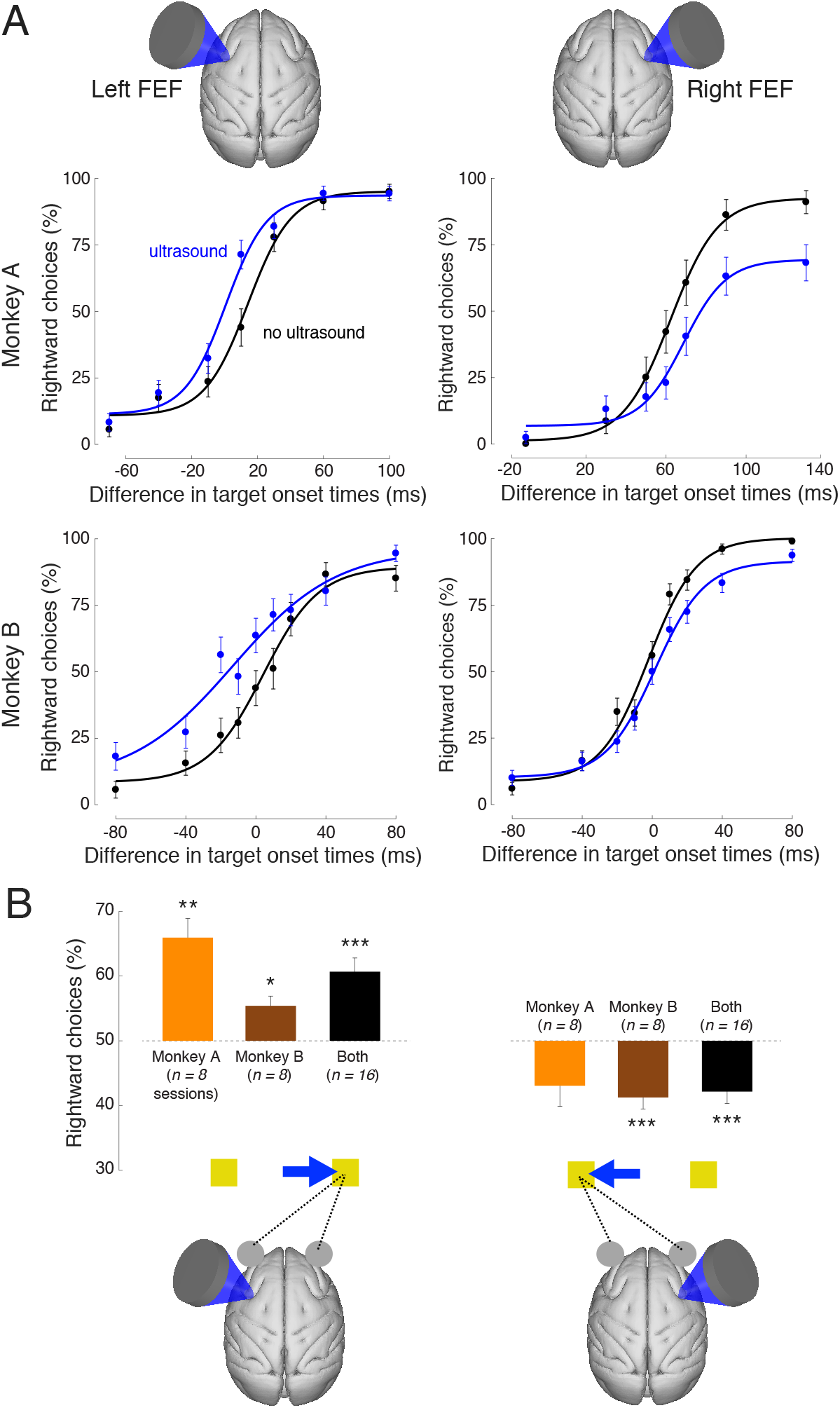
Ultrasonic stimulation of frontal eye fields biases visuo-motor decisions. A) Single session examples. Mean (± s.e.m.) proportion of choices of the rightward target as a function of the difference in target onset times. Positive difference stands for the cases in which the rightward target appeared first. The black data points reflect choice behavior in the trials in which the animal was not stimulated, whereas the blue data points represent choice behavior in the stimulated trials. The data were fit with a four-parameter sigmoid function (see Materials and Methods). The data are presented separately for left and right FEF stimulation sessions (left and right columns; see illustration on top), and for monkey A and B (top and bottom row). B) Quantification of the effects for all sessions. The no stimulation data (black) of each session were first fit with a sigmoid function. Using the fit, we identified the time difference on the abscissa for which the animal chooses both targets in equal proportion (see Materials and Methods). At that point, we then assessed the proportion of rightward choices during the stimulated trials. The number of sessions is provided in parentheses. The stars indicate the cases in which the mean effects statistically differ from equal preference (two-sided t-test; *: *p* < 0.05, **: *p* < 0.01, ***: *p* < 0.001). The illustration on the bottom summarizes the polarity of these biasing effects.

To quantify these effects across all sessions, we measured the proportion of rightward choices on ultrasound trials at the time point at which the animals chose both targets at equal proportion when not stimulated (see Materials and Methods). We did this separately for each session and present the average over sessions (Fig. 2B). This analysis shows that targeting left FEF increased the proportion of rightward choices (Fig. 2B, left). The effect was especially strong in monkey A, who chose the rightward target, when stimulated, in 65.9% of cases at the point of otherwise equal preference (50%). That effect was significantly different from 50% across the 8 sessions in this animal (*p* = 0.0015, *t*_7_ = 5.0). The effect is substantial—at a nearly 66% bias, the monkey chose the rightward target nearly twice as often as the leftward target. The effect was statistically significant also in monkey B, and was highly significant across the two animals (*p* = 0.0002, *t*_15_ = 4.9). As in the single session examples (Fig. 2A, right column), the effect reversed polarity on the trials in which right FEF was stimulated (Fig. 2B, right). The monkeys chose the rightward target at the point of equal preference only in 42.2% of cases, and this differed significantly from the 50% equal preference (*p* = 0.0009, *t*_15_ = −4.12). These effects are graphically summarized at the bottom of Fig. 2B. Stimulation of left FEF significantly increased the proportion of rightward choices, whereas stimulation of right FEF significantly increased the proportion of leftward choices. The contralateral nature of these shifts suggests that ultrasound led to a heightened target representation by neurons within the stimulated regions (Fig. 2).

It is worth noting that the point of equal preference for monkey A is generally substantially distinct from 0 (Fig. 2A, top row, black curves). This means that the animal shows an inherent preference for one of the targets. Such an inherent bias is very common in this free choice task, and can vary substantially from session to session (Noudoost and Moore, 2011). To ensure that this variability did not influence our results, monkey B performed the same task with the exception that he was trained to choose the target that appeared first (see Materials and Methods). This step greatly reduces the mean and variance of inherent bias (Kubanek et al., 2015). Fig. 2B shows that from retrospect, this additional control was not necessary—the effects point in the same direction in both animals, and are on average of comparable size.

We investigated the effects on the animals’ decision-making in more detail. In particular, we asked whether ultrasound shifted the decision curves along the horizontal axis and/or changed the curves’ slope (see Materials and Methods). We fitted the decision curves with a two-parameter sigmoid fit (Kubanek et al., 2015), separately for the stimulated and non-stimulated decision curves within each session. We indeed found that ultrasound shifted the curves along the horizontal axis (Fig. 3). Left FEF stimulation shifted the decision curves on average by −7.6 ms (Fig. 3, left), and this effect was highly significant across the sessions (*p* < 0.001, *t*_15_ = −4.3; two-sided t-test). A leftward shift indicates, for a given difference in target onset times, that the animals were more likely to choose the rightward target when stimulated. The effect reversed polarity during right FEF stimulation, shifting the decision curves by +5.6 ms across the sessions. Also this effect was highly significant (*p* = 0.0064, *t*_15_ = 3.2). A rightward shift indicates, for a given difference in target onset times, that the animals were more likely to choose the leftward target when stimulated. These effects were significant within individual sessions (Table 1). Stimulation of left FEF produced a significant horizontal shift of decision curves in 7/16 sessions, by an average of −13.4 ms (compared to −7.6 ms across all 16 sessions). Stimulation of right FEF produced a significant horizontal shift in 3/16 sessions, by an average of +11.1 ms (compared to +5.6 ms across all 16 sessions). These shifts corroborate the finding of Fig. 2 that ultrasound stimulation biased the animals’ choices in the contralateral direction.

**Figure 3.**
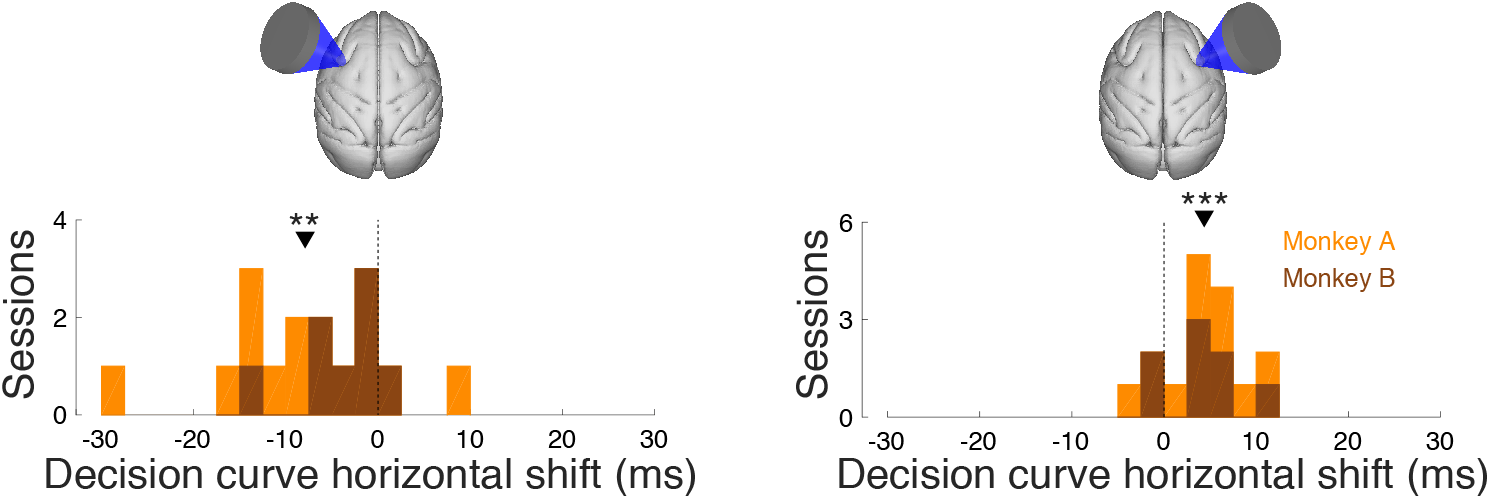
Ultrasound shifts the decision curves to induce contralateral bias in choices. The amount of horizontal shift of the decision curves by ultrasound (blue versus black in Fig. 2A). The data are presented separately for left and right FEF stimulation sessions (left and right panel), and separately for each session in each animal (effect histogram). The stars indicate the effect significance (two-sided t-test; *: *p* < 0.05, **: *p* < 0.01, ***: *p* < 0.001).

**Table 1.**
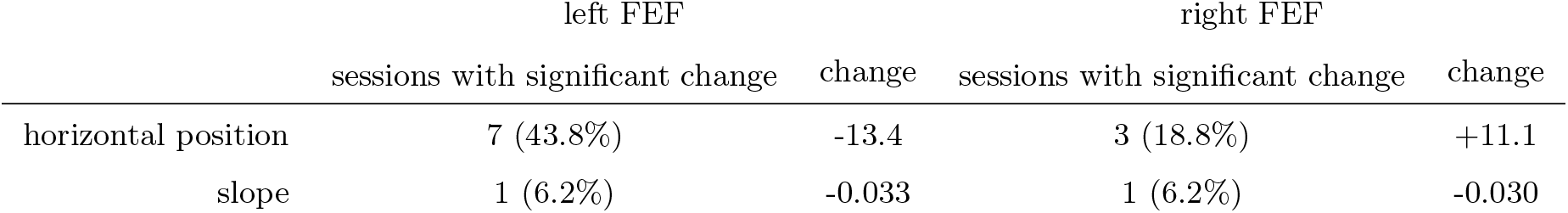
Effects within each session. The table shows the number of sessions (left columns) in which the parameters fitted to each decision curve (rows) changed significantly during ultrasound stimulation. See Materials and Methods for details of the statistical test. The right column shows the average magnitude of the change over the significant sessions. Horizontal position is measured in milliseconds; slope in per millisecond.

Compared to the notable horizontal shifts of the decision curves, ultrasound had only a mild effect on the slope of the curves. Significant shallowing was arguably observed only during left FEF stimulation (a mean change of −0.0095; *p* = 0.032, *t*_15_ = −2.4); not during right FEF stimulation (*p* = 0.22, *t*_15_ = −1.3)). The effect was significant in only 1/16 sessions (Table 1). The lack of substantial shallowing suggests that ultrasound did not notably impair the animals’ ability to distinguish the onsets of the two targets.

Stimulation of the FEF with moderate electric currents is known to elicit saccades into the contralateral hemifield (Bruce et al., 1985; Tehovnik et al., 2000). The magnitude of the stimulating current influences the saccadic endpoint—the larger the current, the farther away from the central fixation point a saccade lands (Bruce et al., 1985; Tehovnik et al., 2000). We therefore tested whether ultrasonic stimulation of FEF produces a similar phenomenon. We found a small but significant effect on horizontal saccadic endpoints (Fig. 4). The effects point in the expected direction based on previous electrical microstimulation studies. For left FEF stimulation, the animals’ saccade endpoints attained an additional 0.20 ° for contralateral saccades (Fig. 4, left). This effect is small given that the eccentricity of the targets was 6 °, but it was highly significant (*p* = 0.001, *t*_14_ = −4.0; two-sided t-test; one session lacked saccade trace data). The effect was not significant for ipsilateral choices (p = 0.95). Conversely, stimulation of right FEF brought the endpoints farther in the opposite, leftward direction (Fig. 4, right). It did so by −0.23 ° (contralateral choices; *p* = 0.039, *t*_15_ = −2.3) and −0.30 ° (ipsilateral choices; *p* = 0.040, *t*_15_ = −2.3). There were no significant effects on vertical saccadic endpoints.. As in a previous study that subjected the FEF to ultrasound of similar parameters (Deffieux et al., 2013), we did not observe ultrasound directly eliciting saccades. We did not observe a significant effect of ultrasound stimulation on the animals’ reaction time, either for contralateral or for ipsilateral choices.

**Figure 4.**
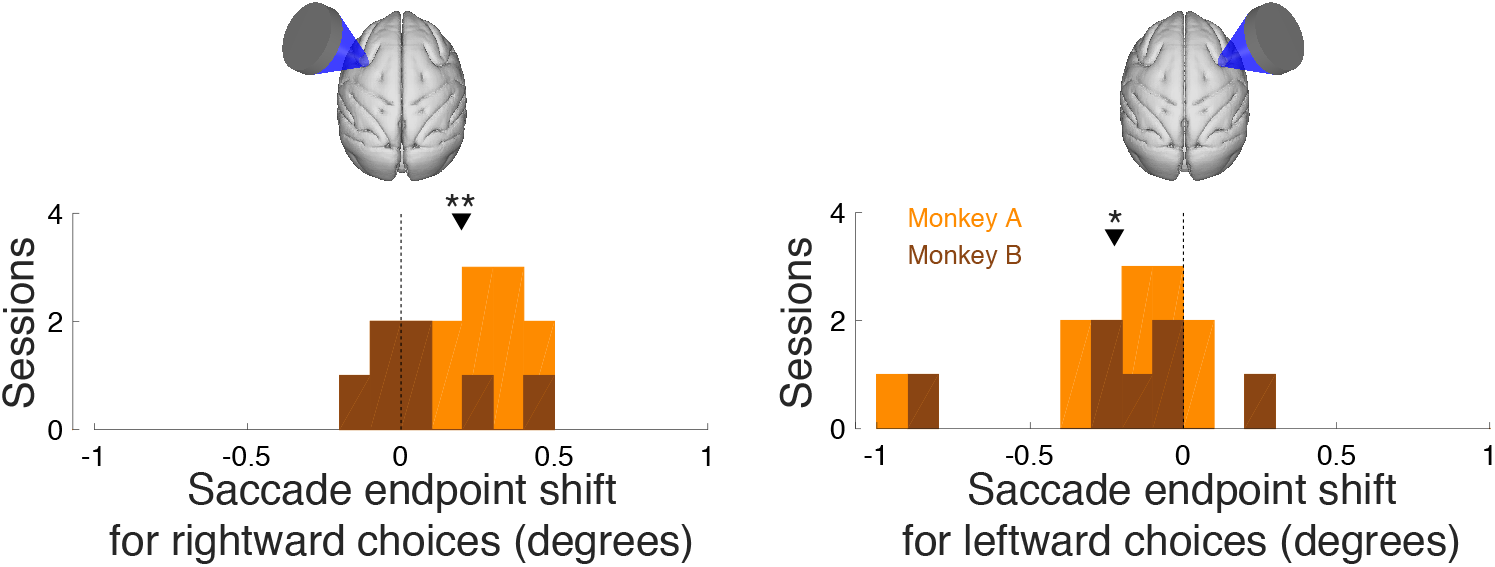
Ultrasound translates saccade endpoints further in the contralateral direction. The histograms show the change in horizontal saccade endpoints (visual degrees) following ultrasonic stimulation as a trial average over each session. The left/right panels specifically show the effects for contralateral (right/left) choices. The stars indicate the effect significance (two-sided t-test; *: *p* < 0.05, **: *p* < 0.01, ***: *p* < 0.001). No significant effects on vertical saccade endpoints were observed.

We investigated how rapidly the effect emerges and whether it is cumulative or, in contrast, whether there is an adaptation. Our task interleaves blocks of stimulated and non-stimulated trials (each 3-6 trials in duration). This block design enables us to assess the dynamics of the ultrasound effects as a function of the number of successively stimulated trials. To do so, we pooled data across right and left FEF stimulation sites and present the average proportion of contralateral choices as a function of trial number within a stimulated and non-stimulated block (Fig. 5). The figure reveals that the biasing effect of ultrasound emerges immediately, on the first stimulated trial within a stimulation block. Interestingly, the effect diminishes in amplitude and becomes insignificant (*p* = 0.082, two-sided t-test) in trial 4 of a block. This suggests that FEF circuitry of the animals adapts to the stimulation, either at the level of FEF neurons or within a broader perceptual system.

**Figure 5.**
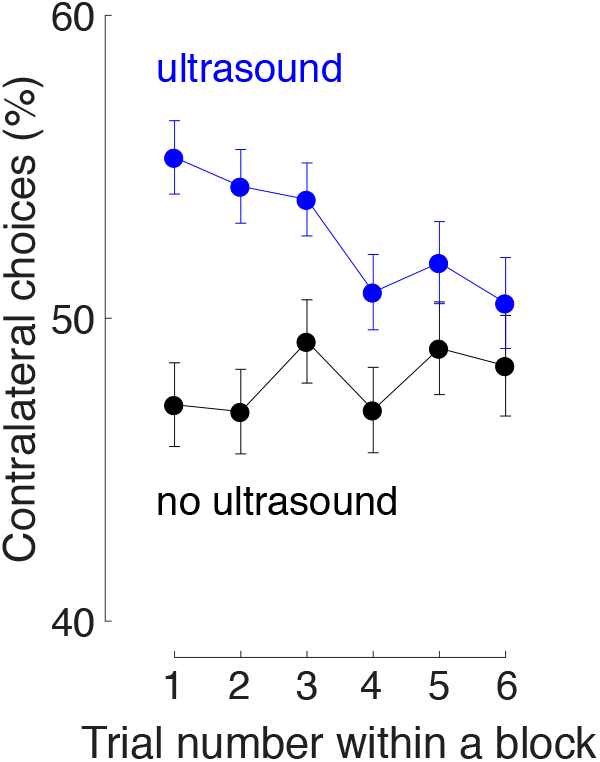
Effect dynamics. Mean ± s.e.m. choices of the contralateral target as a function of trial number within a block, separately for non-stimulated (black) and stimulated (blue) blocks of trials. Data were pooled across left and right FEF stimulation sites.

The differential effects of right and left FEF stimulation (Fig. 2, Fig. 3, Fig. 4) demonstrate that the effects are specific to the stimulated regions. To further validate this observation, we applied the same stimuli to control regions 10 mm more posterior to FEF, i.e., the motor cortex. Motor cortex is not involved in oculomotor choices and so in this case, there should be no effects on animals’ choices or saccade endpoints. We collected data in 11 sessions of left motor cortex stimulation and 11 sessions of right motor cortex stimulation. Besides the change in the stimulation location, data were collected in the same way as with the FEF stimulation. In contrast to FEF stimulation, motor cortex stimulation did not elicit significant biases in choice behavior (Fig. 6). There were no significant shifts from equal preference (left motor cortex: *p* = 0.43, *t*_10_ = −0.82; right motor cortex: *p* = 0.36, t_10_ = 0.95) and no significant horizontal shifts (left motor cortex: *p* = 0.5, *t*_10_ = −0.70; right motor cortex: *p* = 0.31, t_10_ = −1.1). In addition, stimulation of left and right motor cortex did not significantly change the horizontal position or slope of the decision curves within single sessions. Motor cortex stimulation did not produce an effect on horizontal or vertical saccade endpoints (*p* > 0.13 for all four combinations of left and right motor cortex stimulation and contralateral and ipsilateral choices).

**Figure 6.**
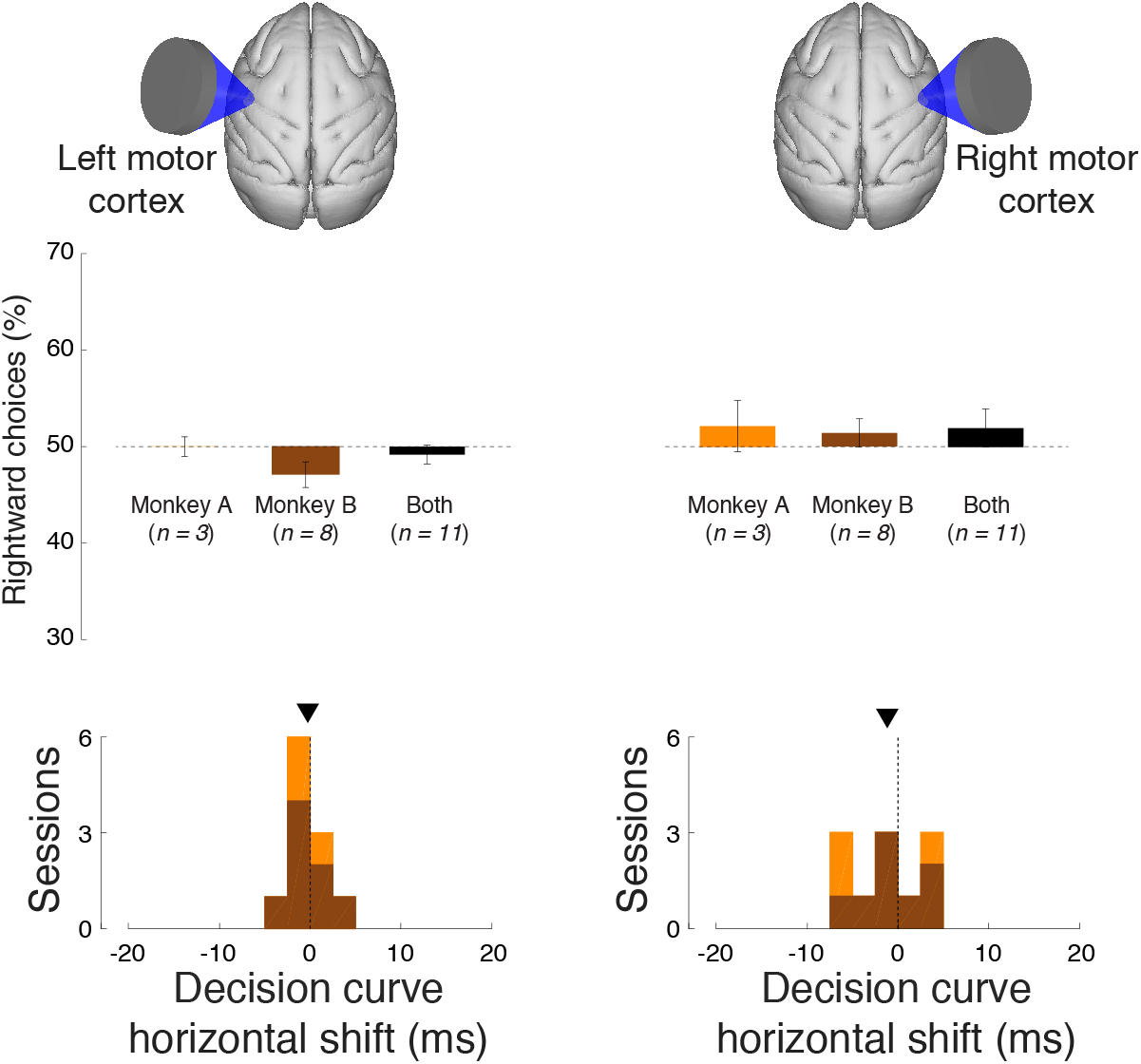
Stimulation of motor cortex had no effect on choice behavior. Same format as in Fig. 2 and Fig. 3, for ultrasound stimulation of left motor cortex (left panel) and right motor cortex (right panel).

## Discussion

We report that transcranial ultrasound can modulate neurons to the extent that it can influence spatial choices of non-human primates. We used an established task that allowed us to assess the polarity, size, and spatial specificity of the effects.

We found that stimulation of left/right FEF biased animals’ choices rightward/leftward (Fig. 2). Based on previous studies, this finding suggests that our pulsed ultrasound stimulus enhanced the representation of the contralateral target within the FEF. In comparison, neuronal inhibition—be it temporary using injected drugs or permanent following lesions—produces shifts of opposite polarity (Rorden et al., 1997; Ro et al., 2001; Schiller and Tehovnik, 2003; Wardak et al., 2004; Kubanek et al., 2015). We therefore conclude that our stimulus, at least in part, excited neurons within the FEF. This finding is supported by a study that recorded neuronal responses within supplementary eye fields in response to ultrasonic FEF stimulation in an anti-saccade task (Wattiez et al., 2017).

The effect was moderate. It was much smaller than effects attained using large injections of potent neuromodulatory drugs (Fig. 1C), but of comparable size to injections of smaller drug volumes (Kubanek et al., 2015) or electrical microstimulation of another node of the oculomotor network, area LIP (Hanks et al., 2006). It is possible that the magnitude of our effects is diminished by our block design, which frequently interleaved stimulated and non-stimulated blocks within a session. In such a block design, carry-over effects from stimulated to non-stimulated trials and/or adaptation to stimulation may reduce the total effect contrast. Nonetheless, the finding that noninvasive ultrasound can produce effects of similar magnitudes as those induced by drugs injected through craniotomies has strong implications for future research of basic brain function. Ultrasound, by virtue of its noninvasiveness and spatial flexibility, may for the first time enable us to screen the contribution of specific brain regions to a given behavior or disease sign systematically, one by one, and in a personalized fashion (Kubanek, 2018).

The ultrasonic effect was specific to the stimulated region, based on two lines of evidence. First, reversing the stimulation hemisphere reversed the effect polarity (Fig. 2, Fig. 3). This constitutes a double dissociation of the effect through different brain regions. In addition, there were no effects when stimulating control regions that are not involved in oculomotor choice, i.e., right and left motor cortex (Fig. 6). Being able to demonstrate a double dissociation regarding the stimulation site is critical to control for generic artifacts that can be associated with propagating ultrasound (Guo et al., 2018; Sato et al., 2018).

We discovered that the biasing effect manifests primarily as a horizontal shift of the decision curves (Fig. 3, Table 1). This finding suggests that our stimulus enhanced the representation of the contralateral target within the FEF. Notably, the slope of the decision curves remained largely intact, with a significant shallowing observed only in 1/16 sessions (Table 1). This indicates that ultrasound did not fundamentally impair the animal’s ability to perceive the stimuli. On the contrary, it apparently enhanced the representation of the contralateral target such that it was more likely to be chosen on a given stimulation trial. Future studies should systematically vary the time and duration within which the ultrasound is applied (we only applied the ultrasound for 300 ms starting 100 ms prior to the appearance of the first target) to specifically impact the sensory, decision-related, and motor stages of the choice process.

There was a small but significant effect on the amplitude of saccades directed into the hemifield contralateral to the stimulated FEF (Fig. 4). Affecting a saccade metric such as saccade endpoint may appear as evidence of ultrasound acting, in part, on motor aspects of saccade planning within FEF. An equally likely possibility, however, is that ultrasound enhanced or shifted the perception of the contralateral target, thus resulting in a slight increase in saccade amplitude. Recording neural activity from FEF and from other, non-stimulated nodes of the oculomotor network such as the parietal area LIP, might help to distinguish between these possibilities.

Our blocked design allowed us to assess the effect progression as a function of the number of consecutive stimuli (Fig. 5). A neuromodulation effect can be constant in time, cumulative—increasing with each additional intervention, or diminishing—decreasing with each additional intervention. We found evidence for the latter kind. The effect emerged immediately, within the first stimulated trial. It then gradually decreased in size until becoming insignificant at about 4-5th consecutive stimulation trial. This indicates that the system adapted to the repetitive stimulation. The nature of this adaptation is currently unknown. One possibility is that the adaptation occurs at the molecular level, whereby the molecular machinery gradually loses sensitivity to repetitive excitation. This possibility is likely given the emerging view that the effects of ultrasound on neurons are of a mechanical kind. In particular, the mechanical forces associated with propagating ultrasound displace membranes and this way open mechanosensitive ion channels (Tyler, 2011; Kubanek et al., 2018; Prieto et al., 2018). It has been demonstrated that such mechanosensing molecules adapt to repetitive mechanical stimulation (Geffeney and Goodman, 2012).

We used a relatively low stimulation frequency (Deffieux et al., 2013) to diminish the role of the skull in neuromodulatory outcomes (Lee et al., 2016). As a consequence, the pressure field associated with our stimulus was relatively broad (Fig. 1D). Although this may appear as a drawback from the perspective of future applications, this in fact provided two benefits in regard to the basic aims of this study. First, the stimulus provided a certain level of tolerance in our FEF targeting. Second, the oblong depth geometry enabled us to stimulate an entire depth of the anterior bank of the arcuate sulcus, a region associated with the FEF. However, it is worth noting that the relatively broad stimulus in part likely influenced other neighboring regions, such as the DLPFC. Future studies can use much more circumscribed stimuli to realize the focusing strength of ultrasound (e.g, about 3 mm half-width when applied through the human skull using large, helmet-like arrays (Ghanouni et al., 2015) and less than 1 mm half-width when applied through a mouse skull (Li et al., 2016)).

The choice paradigm (Fig. 1) used in this study offers several benefits to future studies. First, it can be applied to characterize the effect polarity, size, and spatial specificity of any insonation—and, for that matter—of any neuromodulation protocol. This includes non-invasive (transcranial magnetic/electrical stimulation) and invasive (optogenetics, electrical microstimulation, pharmacological injections) neuromodulation approaches. Second, the block paradigm can be modified (e.g., increased in duration), so that also plastic, long-term effects associated with long-term stimulation (?) can be assessed. Third, the paradigm provides the means to quantify the effect polarity, size, and local specificity noninvasively, from a subject’s choice behavior. And finally, the task is easy to learn and master for animals, which is of tremendous asset in NHP studies.

In summary, we used a task commonly employed in neurology and NHP research to quantify the polarity, size, and spatial specificity of the effects of transcranial focused ultrasound on neurons in NHPs. We demonstrate that ultrasound can noninvasively modulate neurons in oculomotor circuits and so substantially influence animals’ spatial decisions. We show that the effect points in the contralateral direction, that its size is comparable to moderate injections of neuromodulatory drugs into oculomotor regions, and that the effect is localized since stimulating the opposite hemisphere reverses the effect’s polarity. A major contribution of the study is the demonstration that the effects of ultrasound on neurons in NHPs are of sufficient magnitude to modulate behavior. This is critical because flexible, systematic neuromodulation of neural circuits can enable rapid and causal screening of the candidate circuits involved in specific disorders. This way, ultrasonic neuromodulation may realize its potential in noninvasive and personalized diagnoses of a variety of brain conditions, and provide a tool to enable new, causal investigations of basic brain function in humans.

## Materials and Methods

### Subjects

Two adult male rhesus monkeys (macaca mulatta, monkey A: 13 kg, monkey B: 7 kg) participated in this study. The animals sat head-fixed in a custom designed monkey chair in a completely dark room. Visual stimuli were displayed on a LCD monitor positioned 25 cm in front of the animals’ eyes. Eye position was monitored using a camera (EyeLink). All procedures conformed to the Guide for the Care and Use of Laboratory Animals and were approved Stanford University Institutional Animal Care and Use Committee.

### Task

Monkeys were trained in a visual discrimination task that has been used to measure neural deficits or enhancements in previous studies (Rorden et al., 1997; Schiller and Chou, 1998; Ro et al., 2001; Wardak et al., 2002; Scherberger et al., 2003; Balan and Gottlieb, 2009). Briefly, in this task, monkeys first acquired a fixation target. After a short delay, a first target (gray square of 0.5 ° by 0.5 °) appeared in the left (right) part of the screen, 6 ° away from the center of fixation. After a random delay ([0, 130] ms, adjusted to the performance of each monkey), a second target, of identical parameters, appeared in the right (left) part of the screen. The order of appearance (left versus right) was randomized from trial to trial. Once presented, both targets remained present until a choice was made. To receive a liquid reward, the animals had to make a saccade to one of the targets within 1 s after the appearance of the first target. The animal had to make the saccade within a 2 ° acceptance window and remain in the window for at least 100 ms. In monkey A, choice of either target was rewarded. This free choice task is commonly associated with a substantial bias preference for one of the targets, and this bias varies considerably across days (Noudoost and Moore, 2011). To test whether our results are independent of this bias, monkey B was only rewarded for choosing the target that appeared first. This effectively mitigates a bias (Kubanek et al., 2015). The effects of ultrasonic stimulation had the same polarity and were comparable in magnitude in both monkeys.

### Ultrasonic stimulation

Ultrasound was applied in blocks of 3-6 trials (the specific number was drawn from uniform distribution bounded between 3 and 6) and was strictly interleaved with blocks of 3-6 trials in which ultrasound was not applied. In monkey A (B), ultrasound was applied on average in 379 (481) trials per session; the stimulated trials constitute 50% of total trials. We stimulated the macaque frontal eye fields (FEF) using the same approach and transducer as described previously (Deffieux et al., 2013). The main difference is that our transducer operated at 270 kHz instead of 320 kHz, and we used a longer stimulus, 300 ms instead of 100 ms. The single element transducer (H-115, diameter 64 mm, Sonic Concepts), geometrically focused to 63 mm, was used with a coupling cone filled with agar. The height of the cone was chosen such that the geometric focus was located 5 mm below the skull, to ensure that ultrasound stimulated neurons within the entire depth of the arcuate sulcus. Pulsed stimulus (300 ms duration, 500 Hz pulse repetition frequency, 50% duty cycle; Fig. 1D) was generated using a commercial function generator (33520B, Keysight) and subsequently amplified using a commercial amplifier (A150, E&I). As previously (Deffieux et al., 2013), the output pressure maximum was set to 0.6 MPa. The pressure field (Fig. 1D) was characterized in vitro in free field, using Aims III (Onda) water tank filled with distilled and degassed water. The same coupling cone filled with agar gel as that used in the main experiment was used in these measurements; no ex-vivo skull was present. The measurements were taken using a calibrated fibre-optic hydrophone (Precision Acoustics). The distribution of the pressure field was measured using a robotized moving stage (Aims III) and characterized in 1 (2) mm steps in the lateral (axial) dimensions (Fig. 1D). The animals were not sacrificed following the experiments and so the exact value of the pressure below the skull is not known. During the experiment, the animals’ hair was shaved and degassed ultrasound gel applied on the skin to mediate good acoustic coupling between the agar-filled coupling cone and the skin. The FEF target was localized using anatomical MRI images.

### Characterization of decision curves

We fitted decision curves of each session with a sigmoid function. The fit was performed separately for stimulated and non-stimulated data (e.g., blue and black curves in Fig. 2A). We used the same four-parameter fit as a previous study (Kubanek et al., 2015). This fit is mathematically equivalent to logistic fit, with the exception that it features two additional parameters to capture also vertical properties (vertical scale and vertical position) of the decision curves:

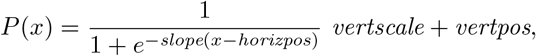

where *P*(*x*) is the probability (frequency) of choosing the rightward target (i.e., the individual points of each decision curve), *slope* defines the steepness of the curve, *horizpos* corresponds to the position of the curve along the horizontal axis (for *vertscale* = 1 and *vertpos* = 0, *x* = horizpos corresponds to the point of equal preference), *vertscale* is a scaling multiplier along the vertical axis, and *vertpos* is a biasing term along the vertical axis.

The parameters were fitted to the choice data using non-linear minimization (function `fminsearch` in Matlab), minimizing the squared error between the fitted and the actual psychometric curves.

From this equation, the point of equal preference used in the analysis of Fig. 2B, *x*_50_ is determined as

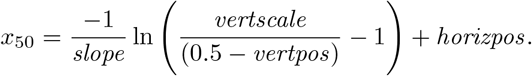

*x*_50_ was computed for each session using the non-stimulated decision curve. Using the stimulated decision curve, we then evaluated the proportion of rightward choices *P*(*x*_50_) at this point of equal preference. Fig. 2B shows the average *P*(*x*_50_) across the individual sessions.

In Fig. 3, “horizontal shift” is the difference in the fitted *horizpos* values of the stimulated and non-stimulated curves.

### Assessment of effects within individual sessions

To assess how the fitted parameters changed between stimulated and non-stimulated trials in each session, we performed a randomization test. In this test, the non-stimulated binary choice data for each difference in target onset times were sampled, with replacement, 10,000 times. Each of these re-sampled decision curves were fit with a sigmoid function. The fitting procedure was the same as above with the exception that we only used *horizpos* and *slope* as parameters. The main conclusions remain the same regardless of whether we use two or four parameters, but using two parameters helped to increase the statistical power of the analysis (we only collected a maximum of 500 stimulated trials per session). The fits produced a null distribution of 10,000 values for each parameter. We then fitted the two parameters to the stimulated curve, and evaluated the probability that the parameters were drawn from the respective null distributions. If the probability was less than 0.01 for a given parameter, Bonferroni-corrected for the number of sessions, the change was taken as significant.

## Acknowledgements

This work was supported by the NIH grant K99NS100986 (JK), Stanford Medicine Dean’s Fellowship (JK), and Howard Hughes Medical Institute (JB, TM, WN). We thank Sania Fong and Shellie Hyde for technical and procedural assistance.

## References

Balan PF, Gottlieb J (2009) Functional significance of nonspatial information in monkey lateral intraparietal area. The Journal of Neuroscience 29:8166–76.

Bruce CJ, Goldberg ME, Bushnell MC, Stanton GB (1985) Primate frontal eye fields. ii. physiological and anatomical correlates of electrically evoked eye movements. Journal of neurophysiology 54:714–734.

Deffieux T, Younan Y, Wattiez N, Tanter M, Pouget P, Aubry JF (2013) Low-intensity focused ultrasound modulates monkey visuomotor behavior. Current Biology 23:2430–2433.

Fini M, Tyler WJ (2017) Transcranial focused ultrasound: a new tool for non-invasive neuromodulation. International Review of Psychiatry 29:168–177.

Fomenko A, Neudorfer C, Dallapiazza RF, Kalia SK, Lozano AM (2018) Low-intensity ultrasound neuromodulation: An overview of mechanisms and emerging human applications. Brain stimulation 11:1209–1217.

Geffeney SL, Goodman MB (2012) How we feel: ion channel partnerships that detect mechanical inputs and give rise to touch and pain perception. Neuron 74:609–619.

Ghanouni P, Pauly KB, Elias WJ, Henderson J, Sheehan J, Monteith S, Wintermark M (2015) Transcranial MRI-guided focused ultrasound: a review of the technologic and neurologic applications. American Journal of Roentgenology 205:150–159.

Guo H, Hamilton II M, Offutt SJ, Gloeckner CD, Li T, Kim Y, Legon W, Alford JK, Lim HH (2018) Ultrasound produces extensive brain activation via a cochlear pathway. Neuron 98:1020–1030.

Hameroff S, Trakas M, Duffield C, Annabi E, Gerace MB, Boyle P, Lucas A, Amos Q, Buadu A, Badal JJ (2013) Transcranial ultrasound (tus) effects on mental states: a pilot study. Brain stimulation 6:409–415.

Hanks TD, Ditterich J, Shadlen MN (2006) Microstimulation of macaque area lip affects decision-making in a motion discrimination task. Nat Neurosci 9:682–9.

Kamimura HA, Wang S, Chen H, Wang Q, Aurup C, Acosta C, Carneiro AA, Konofagou EE (2016) Focused ultrasound neuromodulation of cortical and subcortical brain structures using 1.9 mhz. Medical physics 43:5730–5735.

Kim H, Chiu A, Lee SD, Fischer K, Yoo SS (2014) Focused ultrasound-mediated non-invasive brain stimulation: examination of sonication parameters. Brain stimulation 7:748–756.

King RL, Brown JR, Newsome WT, Pauly KB (2013) Effective parameters for ultrasound-induced in vivo neurostimulation. Ultrasound in Medicine & Biology 39:312–331.

King RL, Brown JR, Pauly KB (2014) Localization of ultrasound-induced in vivo neurostimulation in the mouse model. Ultrasound in medicine & biology 40:1512–1522.

Kubanek J (2018) Neuromodulation with transcranial focused ultrasound. Neurosurgical focus 44:E14.

Kubanek J, Li JM, Snyder LH (2015) Motor role of parietal cortex in a monkey model of hemispatial neglect. Proceedings of the National Academy of Sciences 112:E2067–E2072.

Kubanek J, Shukla P, Das A, Baccus SA, Goodman MB (2018) Ultrasound elicits behavioral responses through mechanical effects on neurons and ion channels in a simple nervous system. Journal of Neuroscience pp. 1458–17.

Lee W, Kim HC, Jung Y, Chung YA, Song IU, Lee JH, Yoo SS (2016) Transcranial focused ultrasound stimulation of human primary visual cortex. Scientific reports 6:34026.

Lee W, Kim H, Jung Y, Song IU, Chung YA, Yoo SS (2015) Image-guided transcranial focused ultrasound stimulates human primary somatosensory cortex. Scientific reports 5:8743.

Lee W, Lee SD, Park MY, Foley L, Purcell-Estabrook E, Kim H, Fischer K, Maeng LS, Yoo SS (2016) Image-guided focused ultrasound-mediated regional brain stimulation in sheep. Ultrasound in Medicine & Biology 42:459–470.

Legon W, Ai L, Bansal P, Mueller JK (2018) Neuromodulation with single-element transcranial focused ultrasound in human thalamus. Human brain mapping 39:1995–2006.

Legon W, Sato TF, Opitz A, Mueller J, Barbour A, Williams A, Tyler WJ (2014) Transcranial focused ultrasound modulates the activity of primary somatosensory cortex in humans. Nature Neuroscience 17:322–329.

Li GF, Zhao HX, Zhou H, Yan F, Wang JY, Xu CX, Wang CZ, Niu LL, Meng L, Wu S et al. (2016) Improved anatomical specificity of non-invasive neuro-stimulation by high frequency (5 mhz) ultrasound. Scientific reports 6:24738.

Mehić E, Xu JM, Caler CJ, Coulson NK, Moritz CT, Mourad PD (2014) Increased anatomical specificity of neuromodulation via modulated focused ultrasound. PLOS ONE 9:e86939.

Naor O, Krupa S, Shoham S (2016) Ultrasonic neuromodulation. Journal of Neural Engineering 13:031003.

Noudoost B, Moore T (2011) Control of visual cortical signals by prefrontal dopamine. Nature 474:372–375.

Oppenheim H (1885) Über eine durch eine klinisch bisher nicht verwerthete Untersuchungsmethode ermittelte Form der Sensibilitätsstörung bei einseitigen Erkrankugen des Großhirns. Neurologisches Centralblatt 4:529–533.

Prieto ML, Firouzi K, Khuri-Yakub BT, Maduke M (2018) Activation of piezo1 but not nav1. 2 channels by ultrasound at 43 mhz. Ultrasound in medicine & biology 44:1217–1232.

Ro T, Rorden C, Driver J, Rafal R (2001) Ipsilesional biases in saccades but not perception after lesions of the human inferior parietal lobule. Journal of Cognitive Neuroscience 13:920–929.

Rorden C, Mattingley JB, Karnath HO, Driver J (1997) Visual extinction and prior entry: Impaired perception of temporal order with intact motion perception after unilateral parietal damage. Neuropsychologia 35:421–433.

Sato T, Shapiro MG, Tsao DY (2018) Ultrasonic neuromodulation causes widespread cortical activation via an indirect auditory mechanism. Neuron 98:1031–1041.

Scherberger H, Goodale MA, Andersen RA (2003) Target selection for reaching and saccades share a similar behavioral reference frame in the macaque. Journal of Neurophysiology 89:1456–1466.

Schiller PH, Chou Ih (1998) The effects of frontal eye field and dorsomedial frontal cortex lesions on visually guided eye movements. Nature neuroscience 1:248–253.

Schiller PH, Tehovnik EJ (2003) Cortical inhibitory circuits in eye-movement generation. European Journal of Neuroscience 18:3127–3133.

Tehovnik EJ, Sommer MA, Chou IH, Slocum WM, Schiller PH (2000) Eye fields in the frontal lobes of primates. Brain Research Reviews 32:413–448.

Tufail Y, Yoshihiro A, Pati S, Li MM, Tyler WJ (2011) Ultrasonic neuromodulation by brain stimulation with transcranial ultrasound. Nature Protocols 6:1453–1470.

Tyler WJ (2011) Noninvasive neuromodulation with ultrasound? a continuum mechanics hypothesis. The Neuroscientist 17:25–36.

Tyler WJ, Lani SW, Hwang GM (2018) Ultrasonic modulation of neural circuit activity. Current opinion in neurobiology 50:222–231.

Wardak C, Olivier E, Duhamel JR (2002) Saccadic target selection deficits after lateral intraparietal area inactivation in monkeys. The Journal of Neuroscience 22:9877–9884.

Wardak C, Olivier E, Duhamel JR (2004) A deficit in covert attention after parietal cortex inactivation in the monkey. Neuron 42:501–508.

Wattiez N, Constans C, Deffieux T, Daye PM, Tanter M, Aubry JF, Pouget P (2017) Transcranial ultrasonic stimulation modulates single-neuron discharge in macaques performing an antisaccade task. Brain stimulation 10:1024–1031.

Ye PP, Brown JR, Pauly KB (2016) Frequency dependence of ultrasound neurostimulation in the mouse brain. Ultrasound in medicine & biology 42:1512–1530.

